# Identifying SARS-CoV-2 Antiviral Compounds by Screening for Small Molecule Inhibitors of nsp5 Main Protease

**DOI:** 10.1101/2021.04.07.438806

**Authors:** Jennifer C Milligan, Theresa U Zeisner, George Papageorgiou, Dhira Joshi, Christelle Soudy, Rachel Ulferts, Mary Wu, Chew Theng Lim, Kang Wei Tan, Florian Weissmann, Berta Canal, Ryo Fujisawa, Tom Deegan, Hema Nagara, Ganka Bineva-Todd, Clovis Basier, Joseph F Curran, Michael Howell, Rupert Beale, Karim Labib, Nicola O’Reilly, John F.X Diffley

## Abstract

The coronavirus 2019 (COVID-19) pandemic, caused by the severe acute respiratory syndrome coronavirus 2 (SARS-CoV-2), spread around the world with unprecedented health and socio-economic effects for the global population. While different vaccines are now being made available, very few antiviral drugs have been approved. The main viral protease (nsp5) of SARS-CoV-2 provides an excellent target for antivirals, due to its essential and conserved function in the viral replication cycle. We have expressed, purified and developed assays for nsp5 protease activity. We screened the nsp5 protease against a custom chemical library of over 5,000 characterised pharmaceuticals. We identified calpain inhibitor I and three different peptidyl fluoromethylketones (FMK) as inhibitors of nsp5 activity *in vitro*, with IC_50_ values in the low micromolar range. By altering the sequence of our peptidomimetic FMK inhibitors to better mimic the substrate sequence of nsp5, we generated an inhibitor with a subnanomolar IC_50_. Calpain inhibitor I inhibited viral infection in monkey-derived Vero E6 cells, with an EC50 in the low micromolar range. The most potent and commercially available peptidyl-FMK compound inhibited viral growth in Vero E6 cells to some extent, while our custom peptidyl FMK inhibitor offered a marked antiviral improvement.

## Introduction

The COVID-19 pandemic, caused by the SARS-CoV-2 virus, emerged late in 2019 and rapidly spread around the world, developing into the worst health crisis of the 21^st^ century [1, 2]. As of February 2021, a year after the WHO declared the outbreak a public health emergency of international concern, over 100 million cases have been confirmed, with more than 2.4 million deaths attributed to COVID-19 [3]. Over 60 different vaccines have now reached the clinical development stage, with several approved worldwide [4]. Vaccinating vulnerable people against SARS-CoV-2 is of paramount importance. However, new variants of the virus are emerging, and it is unclear how long vaccines will remain effective [5, 6]. Thus, to combat this pandemic most effectively, antiviral drugs are needed as complements to vaccines.

SARS-CoV-2 encodes at least nine enzymes that are important for viral proliferation and thus are attractive targets for antiviral drugs. In contrast to the rapidly evolving structural proteins of the virus that the vaccines are based on, these enzymes are highly conserved between different coronaviruses [7, 8]. This suggests that antivirals inhibiting these enzymes may be useful as pan-coronavirus treatments and as the virus becomes resistant to existing vaccines. COVID-19 is already the third zoonotic coronavirus after SARS-CoV-1 and MERS-CoV that emerged as global health threats during the last two decades [9], so the availability of pan-coronavirus treatments as a first line of defence against novel coronaviruses may be crucial in the future.

The first two-thirds of the SARS-CoV-2 genome encodes sixteen non-structural proteins (nsps), which are required for viral proliferation [10, 11]. They are encoded in two large overlapping open reading frames (ORF 1a and ORF 1ab). Upon entry into the host cell, these are translated into two polyproteins (pp1a and pp1ab respectively), which are cleaved by two virus encoded cysteine proteases, generating sixteen functional nsps. The viral papain like protease (PLpro), which is encoded within nsp3, excises nsp1-3 [12]. The main viral protease (nsp5) is a chymotrypsin-related protease that cleaves the polyproteins at eleven sites, releasing nsp4-nsp16 [13–15]. The excised nsps are essential for the assembly of the viral replication transcription complex [11]. Inhibition of nsp5, therefore, blocks the viral replication cycle, making it an attractive antiviral drug target [16]. Nsp5 structure and function is conserved across all coronaviruses, with SARS-CoV-1 and 2 sharing roughly 96% sequence identity with the greatest degree of sequence conservation around the active site [14]. Nsp5 cleaves polyproteins after a glutamine residue, which is the case for the vast majority of coronavirus main proteases but is rare in human proteases [8, 17, 18]. This suggests that antivirals targeting the SARS-CoV-2 main protease may have broad spectrum anti-coronavirus inhibitory effects. As part of a larger project to identify inhibitors of all SARS-CoV-2 enzymes, here we describe a high-throughput drug screen to identify inhibitors of the main viral protease nsp5.

## Results

### Expression and purification of nsp5

It has previously been shown that both C- and N-terminal epitope tags inhibit nsp5 dimerization and activity [17, 19]. Therefore, we optimised a purification method for a full length, untagged nsp5 from *E. coli* based on the SUMO/Ulp1 cleavable tag system (Figure 1A)[20]. Figure 1B shows fractions from the purification steps separated in an SDS-polyacrylamide gel; nsp5 migrates at around 36 kDa. This method resulted in a high yield of relatively pure nsp5.

**Figure 1:**
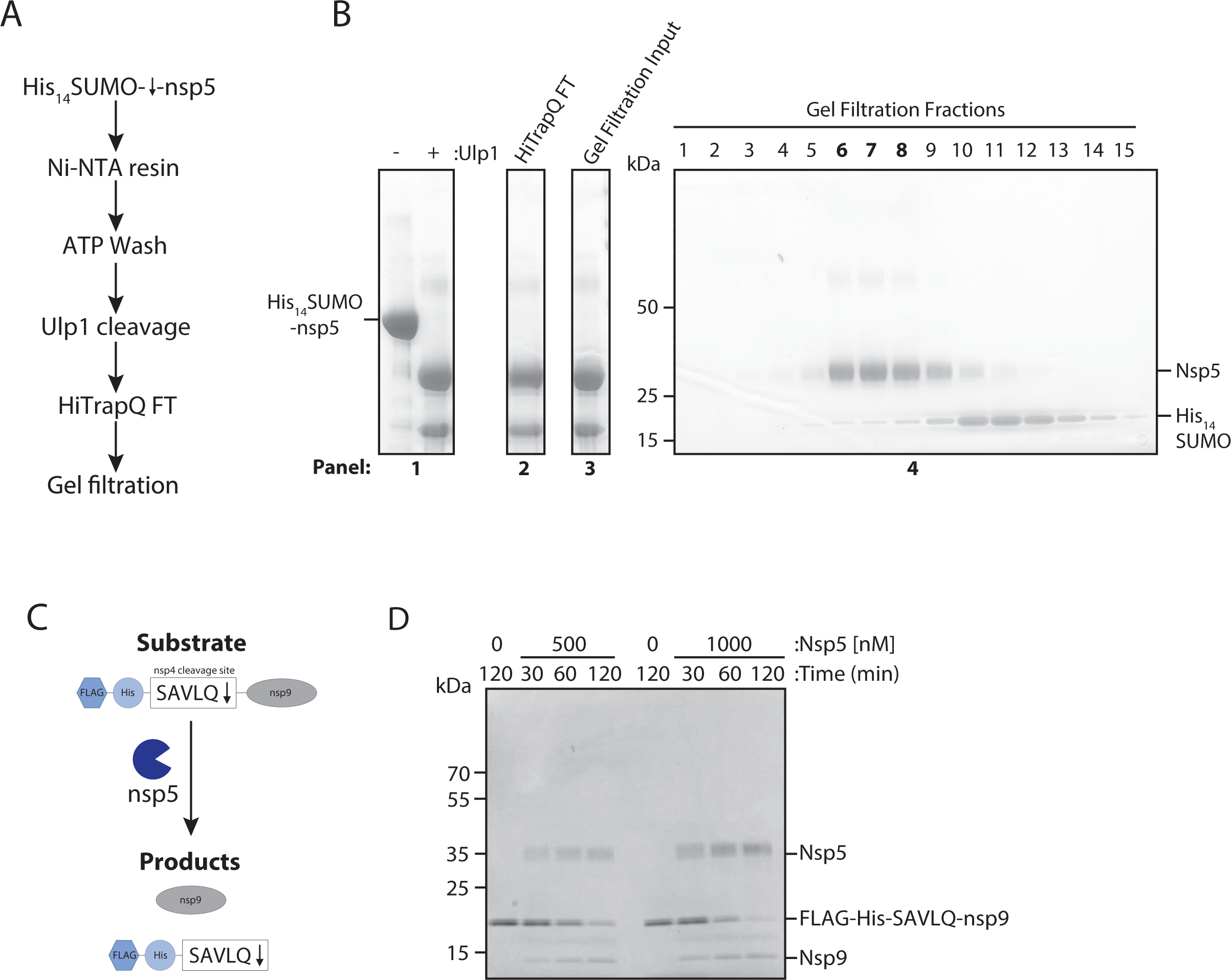
Nsp5 expression, purification and activity test by nsp9 gel-based assay. A) Schematic overview of nsp5 purification. Crude cell extracts containing His_14_-SUMO tagged nsp5 is bound to a Ni-NTA resin, washed with ATP and then eluted by His_14_-SUMO cleavage by the protease Ulp1. Major contaminates are then removed over an anion exchange and gel filtration column. B) Coomassie gel of His_14_-SUMO cleavage. Panel 1: Elution from Ni-NTA beads before and after cleavage of nsp5 by the Ulp1 SUMO-dependent protease. Panel 2: Sample of the flow through from the anion exchange column. Panel 3: Input for gel filtration column. Panel 4: Fractions taken across the major peak of the gel filtration elution. Lanes six through eight (in bold) were pooled and concentrated for use in the HTS screen. C) Schematic overview of nsp9 gel-based cleavage assay. The substrate of FLAG-His- SAVLQ-nsp9 is incubated nsp5. Cleavage of the substrate results in two products: nsp9 and FLAG-His-SAVLQ. D) Gel-based assay for nsp9 cleavage by nsp5 over time at 500 and 1000 nM of nsp5. Nsp5 is visible at around 35 kDa and the substrate of uncleaved nsp9 visible at around 12 kDa.

To examine nsp5 protease activity, we designed a nsp9 gel-based cleavage assay. For this we synthesised a fusion protein substrate with a short linker sequence based on the nsp4- nsp5 junction (SAVLQ) in-between a FLAG-His epitope tag and the nsp9 protein (Figure 1C). We selected the nsp4-5 cleavage site, as it is a natural cleavage site of nsp5 predicted to have the highest affinity [17]. Cleavage of the substrate, which migrates at approximately 20 kDa, by nsp5 results in the release of untagged nsp9, which migrates at approximately 12 kDa. Cleavage of the Flag-His-SAVLQ-nsp9 substrate over time by nsp5 can be observed from as early as 30 minutes at both 0.5 μM and 1 μM of nsp5 (Figure 1D).

### FRET-based primary screen optimisation

To carry out enzymatic characterisation and high throughput screening (HTS) for nsp5 inhibitors, we optimised a Förster (fluorescence) resonance energy transfer (FRET)-based assay (Figure 2A). We used a 10 amino acid-long peptide substrate based on the natural nsp4-nsp5 cleavage site. The peptide is covalently attached to the fluorophore 2- aminobenzoyl (Abz) at its N-terminus and a quencher (N-Tyrosine) at its C-terminus (Figure 2A). Cleavage of the substrate releases the fluorophore from the proximity of the quencher, resulting in an increase of fluorescent signal. Using this substrate, we determined the K_M_ of nsp5 for this substrate to be 33.7 ± 4.7 μM (Figure 2B). This is comparable to a previously reported value of 28.2 μM using a similar FRET substrate [21].

**Figure 2:**
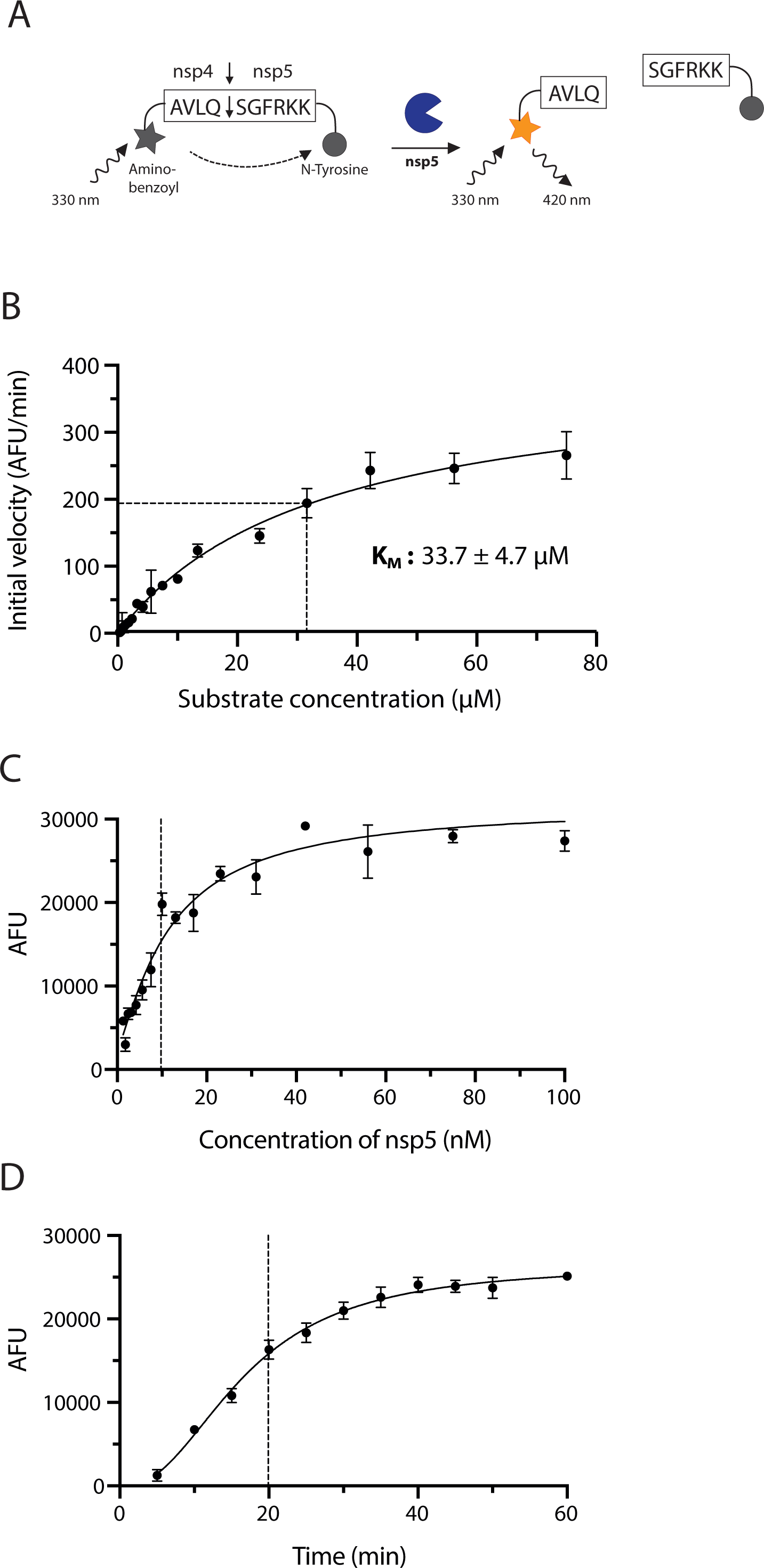
Enzyme assay design and characterisation. A) Schematic of the FRET-based protease assay. In the uncleaved state the fluorescence of aminobenzoyl is quenched by N-tyrosine. Once the protease cleaves the peptide, the fluorescent signal increases proportionally to protease activity. B) Determination of enzyme kinetics. Initial reaction rates over a range of concentrations were plotted against substrate concentration to obtain values for K_M_ (33.72 ± 4.68 μM). Data is plotted as the mean and standard deviation (SD) of 3 replicates. C) Protease activity of nsp5. Increase in fluorescence 20 minutes after the start of the reaction is shown over increasing nsp5 concentrations. Experiment was done in triplicate. D) Time course of the nsp5 hydrolysis reaction. Increase in fluorescence at 10 μM enzyme concentration over time is shown. Experiment was done in triplicate.

We found that, at an enzyme concentration of 10 nM, the reaction is roughly linear for the first 20 minutes at room temperature (19 – 23 °C) (Figure 2C-D). We used a substrate concentration of 20 μM for the HTS, which is relatively close to the K_M_ value to ensure that competitive as well as non-competitive inhibitors can be identified [22]. We also found that buffer conditions had a profound effect on activity, especially when using automated liquid handling (Figure S1). In brief, it proved essential to include glycerol (between 5-10%) and detergent (e.g. 0.01-0.02% Tween 20) to stabilise the enzyme. Inclusion of detergent also prevents compound aggregation ([23] and Zeng *et al.* this issue).

### High throughput screen hit identification and validation

We carried out a high throughput screen (HTS) of a custom library with over 5,000 compounds at two drug concentrations of 4 μM and 0.8 μM. For the HTS, we incubated the compounds with nsp5 before reaction initiation to allow for the detection of slow-binding inhibitors. We defined primary hits as compounds that reduced nsp5 activity by more than 30% at the higher compound concentration (Figure 3A, see Methods). We identified 27 primary hits that met this criterion and ranked them by percentage inhibition of nsp5 activity at both drug concentrations (Table 1) and validated them according to the scheme presented in Figure 3B.

**Figure 3:**
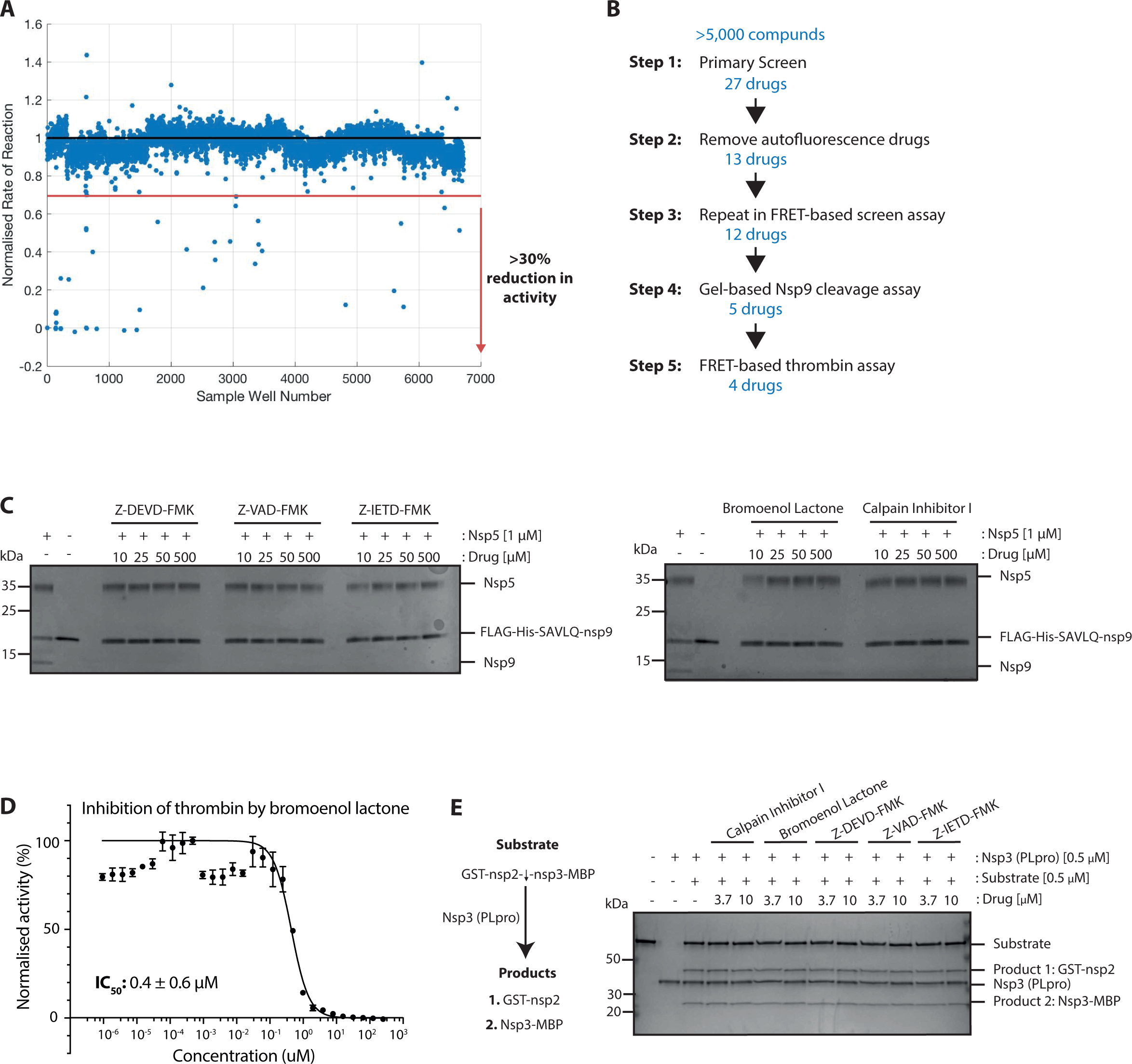
Primary screen hit identification and validation. A) Scatterplot of all screen results at the low drug concentration (0.8 μM). The red dotted line indicates the threshold of a 30% reduction in enzyme activity required to class as an initial hit. All points below the red line were considered in the initial hit list. B) Schematic overview of validation steps and number of hits at each stage. C) Gel-based nsp9 cleavage assays of initial hits from the HTS. Only drugs shown were those that inhibited nsp5 activity at any of the indicated concentrations. D) A half-maximum inhibitory concentration (IC_50_) value for bromoenol lactone against nsp5 activity was determined by non-linear regression (shown in black). Data is plotted as mean and standard deviation (SD) of 3 replicates. E) Gel-based assay of substrate cleavage by the other SARS-CoV-2 protease PLpro (nsp3). The five hits from the screen were tested for specificity against PLpro protease.

**Table 1:**
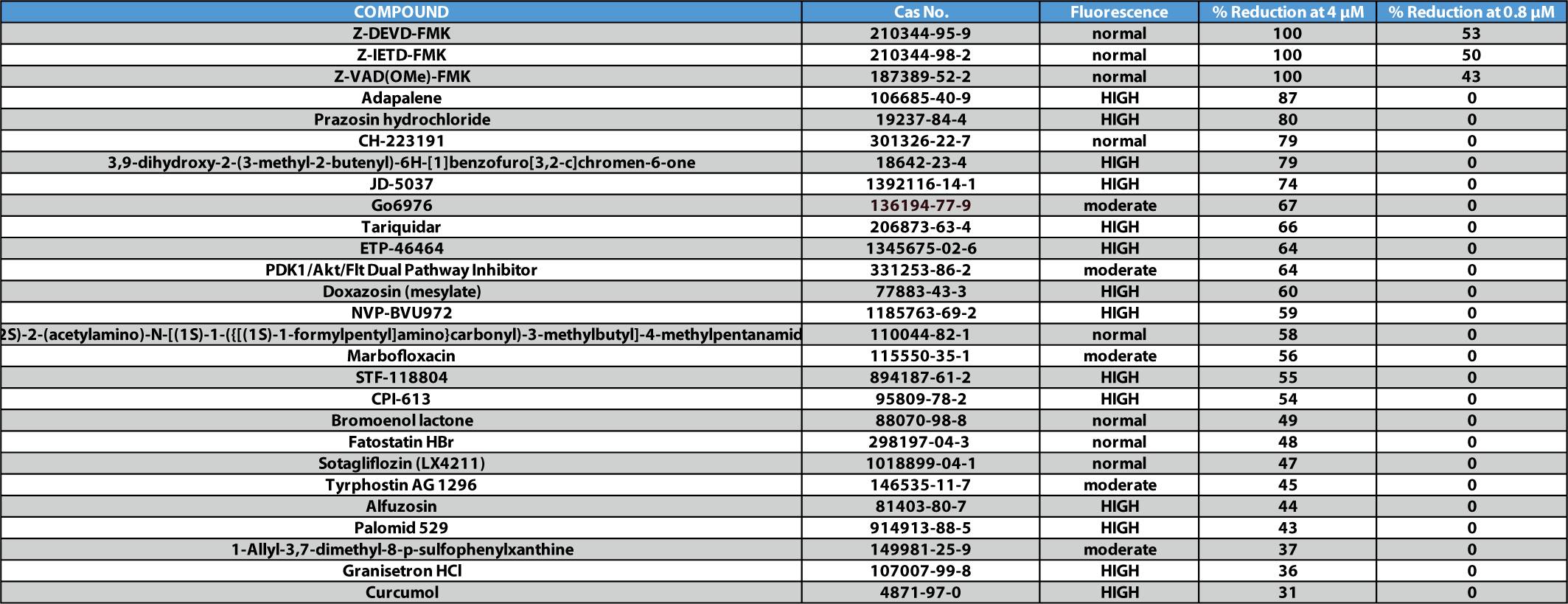
Full primary hit list. Table showing ranked initial hits with compound name, fluorescence reading type and percentage inhibition at both primary screen concentrations (4 and 0.8 μM). Those with “HIGH” fluorescence readings were either reading at over the detectable range in the screen wavelength or completely saturated. Such a reading would result in a decreased rate in reaction due to how reaction rates were determined via MATLAB. Those reading at “moderate’ had slightly increased fluorescence at both screen concentrations, but a rate of reaction could still be determined. Finally, those with “normal” fluorescence readings were drugs that did not interfere with the fluorescence signal at the primary screen emission wavelength at all.

Some compounds within the library interfered with the HTS emission wavelength (420 nm), increasing the fluorescence of the first time point and subsequent time points. Due to a maximum range of fluorescence detection these reactions became saturated over the course of the reaction resulting in an artificially reduced rate of reaction being determined for these compounds. We classed drug compounds that did not interfere with the HTS emission wavelength as “normal”. Those that interfered slightly but a reduction in nsp5 activity was still evident we classed as “moderate”. Finally, those that greatly interfered with the HTS fluorescence and so were likely artificial hits we classed as “high” (Table 1). We excluded fourteen drugs from the primary hit list, which showed “high” autofluorescence. We were unable to source one of the compounds, so we tested twelve compounds further as nsp5 inhibitors.

We first re-tested these twelve compounds in the FRET-based assay over a range of drug concentrations from 0 to 500 μM and calculated IC_50_ values where possible. Four compounds did not show reproducible inhibition of nsp5 across any of the tested drug concentrations (Figure S2A) and we, therefore, discarded them. Tryphostin artificially increased fluorescence in the FRET- based assay at some concentrations tested, reducing the number of points available for IC_50_ calculation. Therefore, we also tested it in the nsp9 gel-based cleavage assay (Figure S2A). As it did not inhibit in either validation step, we discarded it. Fatostatin HBr and PDK1 /Akt /Flt Dual pathway showed some inhibition of nsp5 in the FRET-based assay. However, they had relatively high IC_50_ values for nsp5 inhibition and they did not show inhibition at any of the drug concentrations in the nsp9 gel-based assay, thus, we removed them from the final hit list (Figure S2B).

The remaining five drugs inhibited nsp5 in both the FRET-based and gel-based assays: Calpain Inhibitor I (IC_50_: 5.17 ± 0.17 μM), bromoenol lactone (IC_50_: 3.4. ± 0.09 μM), Z-DEVD-FMK (IC_50_: 0.23 ± 0.03 μM), Z-IETD-FMK (IC_50_: 0.23 ± 0.01 μM) and Z-VAD-FMK (IC_50_: 0.16 ± 0.01 μM) (Figure 3C, Figure S2C-D). Calpain Inhibitor I and Z-DEVD-FMK were described recently as nsp5 inhibitors *in vitro* [21, 24]. To test the specificity of the final five drugs we tested them against a different protease, thrombin. We used the same buffer as in the HTS, which contained a non-ionic detergent, which reduces the likelihood of unspecific inhibition via colloidal aggregation [23]. We used a substrate based on the natural thrombin substrate: Aminobenzoyl- LGARGHRPYD-N-Tyrosine [25]. Of the five drugs, only bromoenol lactone inhibited thrombin activity (Figure 3D) indicating that it is not a specific inhibitor of nsp5. Lastly, we tested whether these compounds could inhibit the other viral coronavirus protease (nsp3 PLpro), which is also a cysteine protease [26]. The substrate and assay conditions used in this experiment are described elsewhere (see Lim et al, Biochemical J, this issue).

None of the remaining hits from the primary screen inhibited PLpro, indicating a specificity for nsp5 (Figure 3E). All together, we have identified four specific inhibitors of nsp5 *in vitro* (Table 2).

**Table 2:**
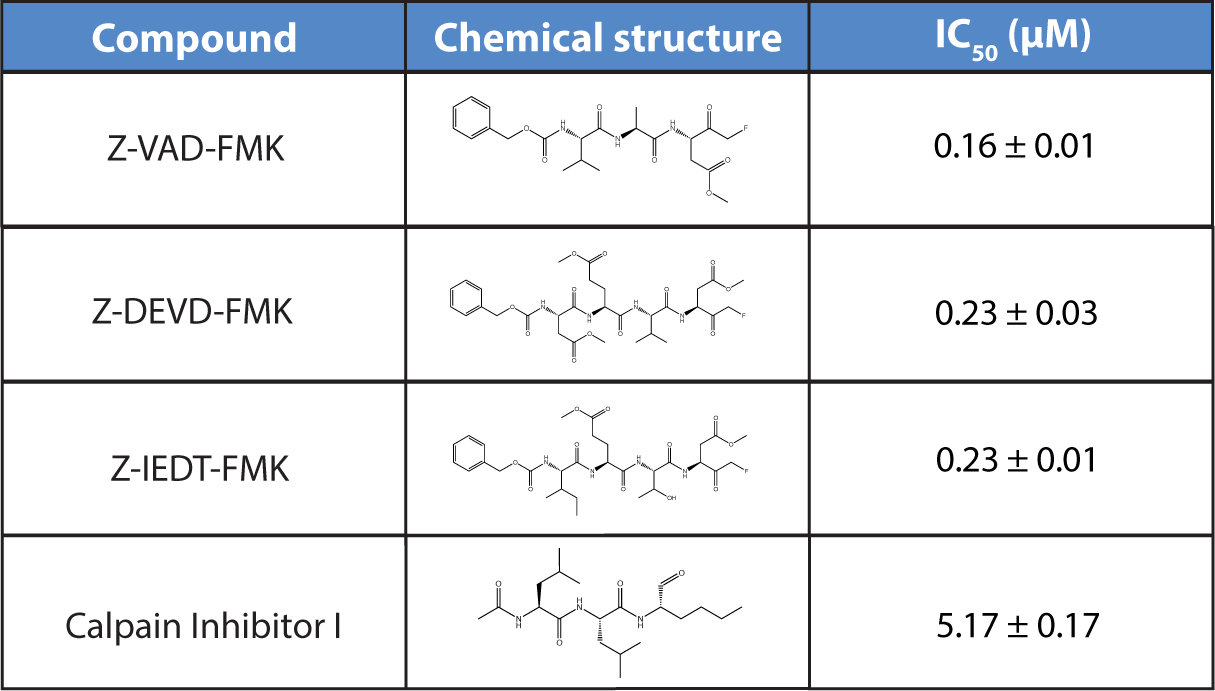
Final hit list. Table showing the final four hits from the HTS. The chemical name, structure and IC_50_ values as determined by non-linear regression are shown.

| Compound | Chemical structure | IC<sub>50</sub> (μM) |
| --- | --- | --- |
| Z-VAD-FMK |  | 0.16 ± 0.01 |
| Z-DEVD-FMK |  | 0.23 ± 0.03 |
| Z-IEDT-FMK |  | 0.23 ± 0.01 |
| Calpain Inhibitor I |  | 5.17 ± 0.17 |

### Evaluation of Ebselen as a nsp5 inhibitor

Ebselen has recently been described as an nsp5 inhibitor [14], but was not identified as an hit in our screen. We wondered whether this discrepancy was due to the presence of reducing agent in our reaction buffers, which was not present in the previous work [14]. We tested Ebselen against nsp5 in the FRET-based assay over a wide range of concentrations in the presence of either DTT or a more physiologically relevant reducing agent, L-Glutathione (GSH) (Figure 4A) [27, 28]. Ebselen inhibited nsp5 only in the absence of a reducing agent with an IC_50_ in the micromolar range. In the presence of either reducing agent there was only a slight reduction of nsp5 activity, at concentrations higher than 100 μM. (Figure 4A). In contrast, Calpain Inhibitor I inhibited nsp5 in the absence of reducing agent as well as the presence of either reducing agent (Figure 4B). The presence of reducing agents did not affect the activity of nsp5 in the absence of inhibitor (Figure S3A). Taken together, these results indicate that Ebselen can only inhibit nsp5 under non-reducing conditions. A similar conclusion was also reached recently by Ma et al. [29].

**Figure 4:**
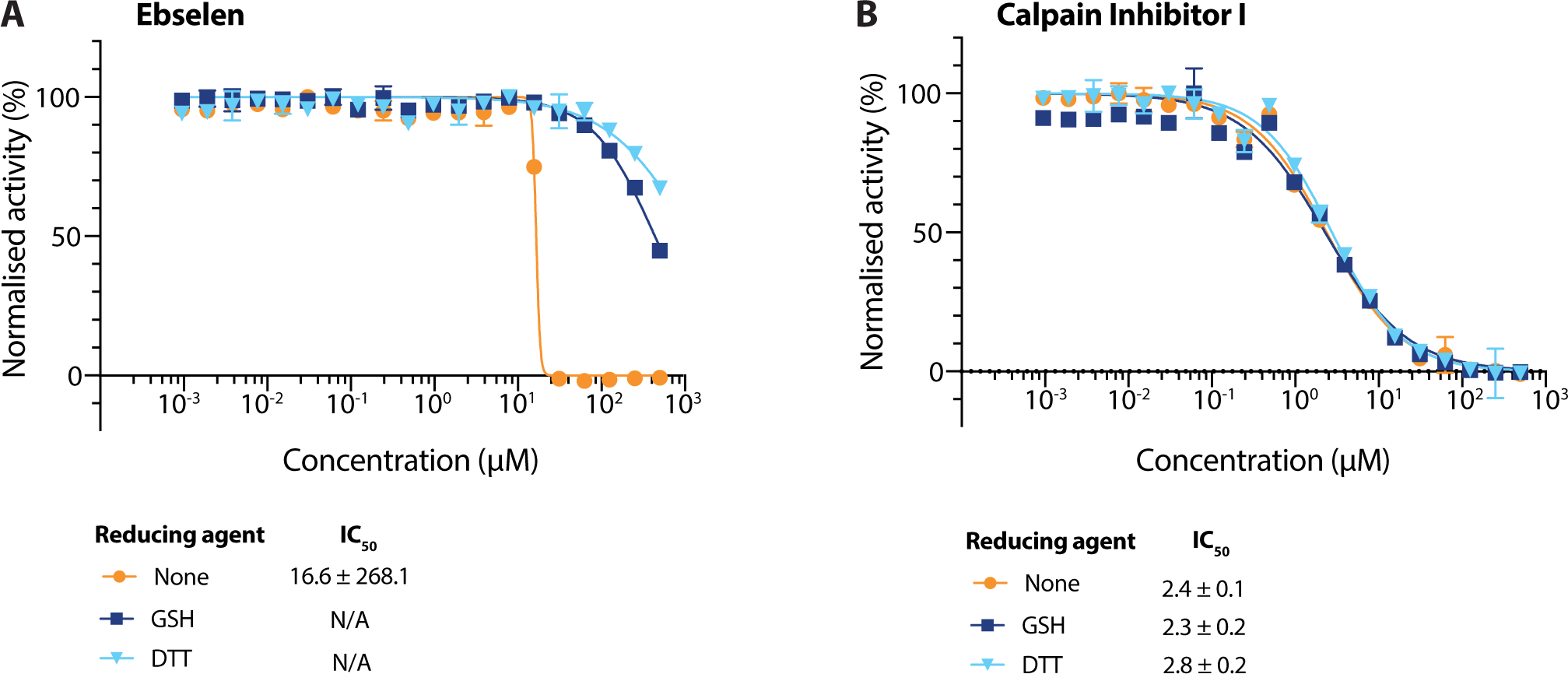
Evaluation of Ebselen as an nsp5 inhibitor. A-B) Dose response curves of Ebselen (A) and Calpain Inhibitor I (B) using different reducing agents in the buffer (DTT, GSH, no reducing agent). IC_50_ values were determined by non-linear regression. Data is plotted as mean and standard deviation (SD), n=3.

### Antiviral activity of drugs in cell-based assay

We next tested the ability of our best hit compounds to inhibit SARS-CoV-2 replication in monkey-derived Vero E6 cells as described by Zeng et al (see this issue) (Figure 5A). Calpain inhibitor I, which has previously been tested for nsp5 inhibition *in vitro* but not in a cell-based assay [21], was the most potent inhibitor of viral infection with an EC50 value of 0.28 ± 0.01 μM. Despite being a good inhibitor *in vitro*, the Z-VAD-FMK inhibitor displayed inhibition only at concentrations greater than 100 μM (Figure 5B-C).

**Figure 5:**
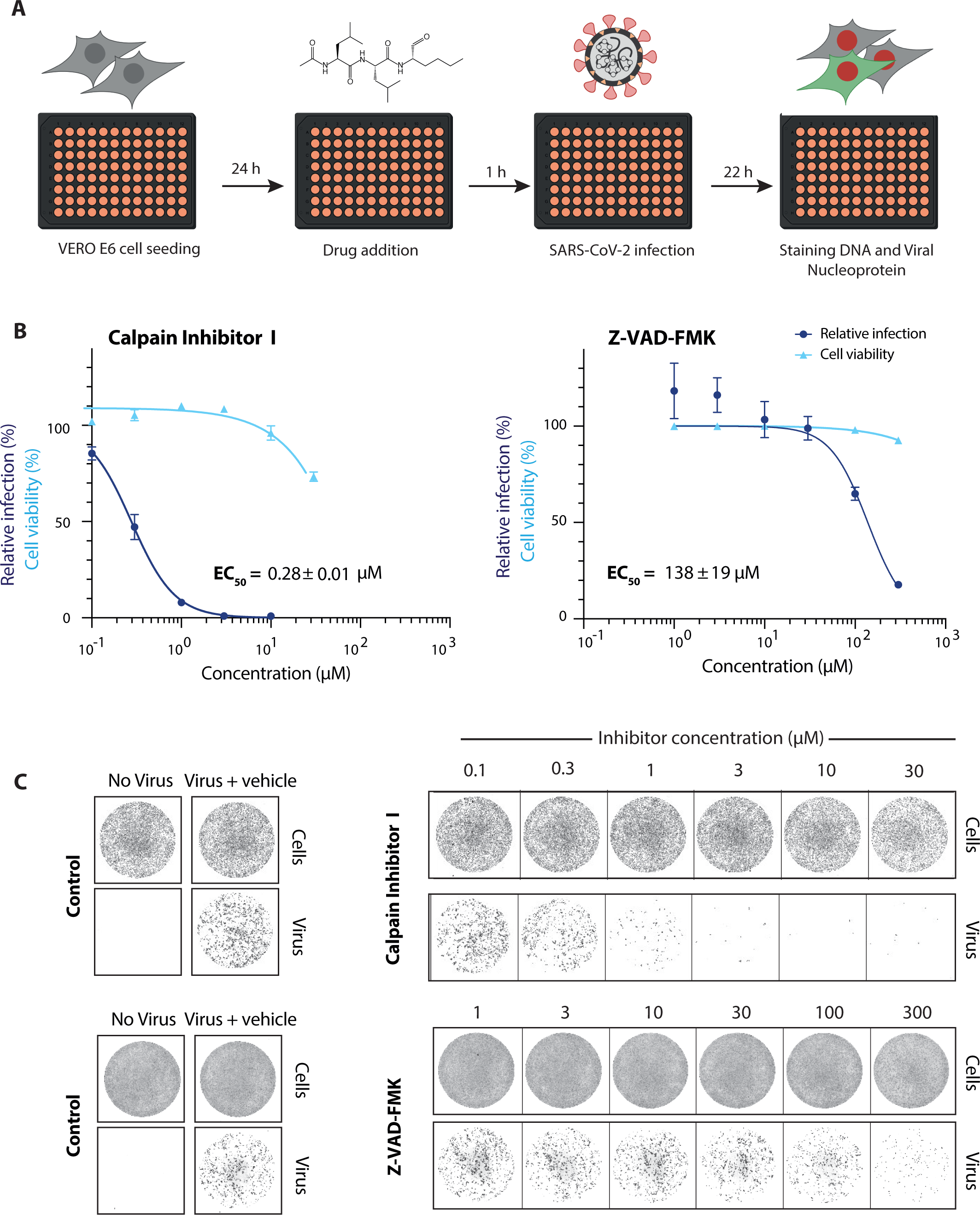
Antiviral activity of selected hits against SARS-CoV-2 in Vero E6 cells. A) Schematic overview of the viral infectivity assay in Vero E6 cells. Vero E6 cells were seeded in a 96-well format and treated with compounds at the indicated concentrations. This was followed by infection with SARS-CoV-2 isolate at a MOI of 0.5 PFU/cell. Afterwards, cells were fixed, stained for DNA with Draq7 and SARS-CoV-2 viral nucleocapsid protein (N protein) with an Alexa-488 labelled antibody and imaged (see Zeng et al. this issue). B) Dose response curves for the validated hits (Calpain Inhibitor I and Z-VAD-FMK) in a viral infectivity assay in Vero E6 cells. Cell viability values represent the area of cells stained with DRAQ7 DNA dye. Viral infection values were measured as the area of viral plaques visualised by immunofluorescent staining of viral N protein. Data is normalised to DMSO only treated control wells (100 %) and plotted as mean and standard deviation (SD), n=3. Half-maximal effective concentration (EC50) values were determined by non-linear regression. C) Representative images for the SARS-CoV-2 viral infectivity assay in Vero E6 cells, for Calpain Inhibitor I and Z-VAD-FMK. Representative wells show Vero E6 cells stained for DNA using DRAQ7 (top panel, labelled cells), and viral N protein immunofluorescence (lower panel, labelled virus).

Combining two antiviral drugs with two different modes of action is a common strategy in treating viral infections as it can increase each drug’s effectiveness and prevent the emergence of drug resistance [30]. We therefore tested the inhibitory capacity of our best hits in the same cell-based assay in combination with remdesivir. Remdesivir is a broad spectrum nucleoside analogue that is currently the only approved antiviral drug against SARS-CoV-2 and targets the RNA-dependent RNA polymerase [31]. None of the nsp5 inhibitors tested showed synergistic or additive effect with remdesivir (Figure S4A-B).

### Improved nsp5 peptidomimetic inhibitors

The three most potent inhibitors *in vitro* were the FMK peptidomimetic inhibitors, originally designed to target caspases [32]. These FMK peptidomimetic inhibitors comprise an N-terminal group that increases cell permeability, a short peptidyl targeting sequence, and a C-terminal functional group which can covalently bind to and inactivate the active cysteine residue in proteases (Figure 6C). We initially tried to investigate the effect of different C-terminal functional groups on inhibitor potency. The functional group in all three of the peptidomimetic inhibitors picked up in our screen is a fluoromethylketone group. While these are non-toxic to cells in culture, experiments in mice showed that their metabolic conversion leads to the production of toxic fluoroacetate [32, 33]. Some safer alternatives of the FMK peptidomimetic inhibitors are available commercially. One such alternative has been developed based on the difluorophenoxymethylketone (OPh) functional group [32]. While they were not present in our drug library, three of them are available commercially; Q-VD-OPh, Q-DEVD-OPh and Q-IETD-OPh. We tested these three drugs in the nsp9 gel-based cleavage assay. Of these, Q-IETD-OPh showed the best nsp5 inhibition in the gel-based assay, Q-DEVD-OPh only displayed inhibition at the highest concentration of drug tested and Q-VD-OPh did not inhibit nsp5 activity (Figure 6A). Similarly, in the FRET-based assay, Q-IETD-OPh was the most effective nsp5 inhibitor, with an IC_50_ value of 17.4 ± 1 μM (Figure 6B). This comparison between peptides with the same amino acid sequence, but differing functional groups, shows that the FMK peptidomimetic inhibitors are more potent inhibitors of nsp5 *in vitro* than the OPh variants.

**Figure 6:**
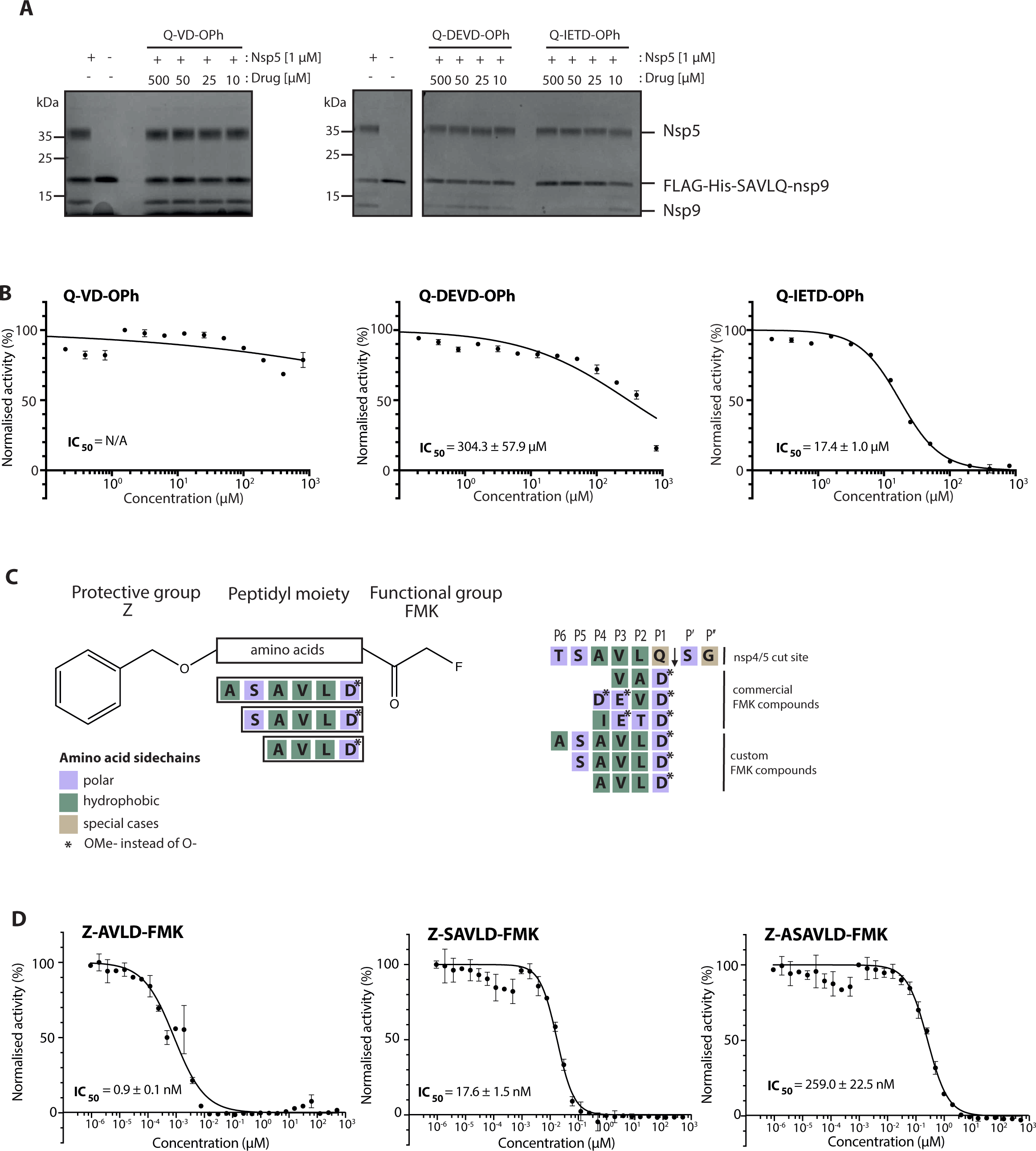
Development of improved nsp5 peptidomimetic inhibitors. A) Gel-based assay of nsp9 cleavage by nsp5 in the presence of the OPh peptidomimetic inhibitors at indicated concentrations. B) Dose response curves of OPh inhibitors. *In vitro* inhibition of nsp5 was measured over a wide range of concentrations of the three OPh inhibitors using the FRET-based assay. A dose response curve for its IC_50_ value was determined by non-linear regression. All data are shown as mean ± SEM, n=3, error bars represent SD. C) Schematic of the FMK peptide inhibitors. Amino acids with different functional side chains are colour coded. D) *In vitro* activity of nsp5 was measured over a wide range of concentrations for the three custom FMK peptides using the FRET-based assay. Dose response curves for IC_50_ values were determined by non-linear regression. All data are shown as mean ± SEM, n=3, error bars represent SD.

Next, we tested whether changing the sequence of the peptidyl moiety would improve the inhibitor potency of the FMK compounds. The three FMK inhibitors identified in the screen mimic the substrate sequence of different caspases [32]. All of them have a common length of three or four amino acids, two hydrophobic residues as well as one charged residue. Work on SARS-CoV-1 nsp5 indicates that bulky hydrophobic residues are key for substrate recognition, especially in the P2 position (Figure 6C) [34]. We hypothesised that altering the amino acid targeting sequence to further mimic an nsp5 substrate might improve the inhibitory effect on nsp5. We synthesised three custom FMK inhibitors, with lengths varying from four to six amino acids (Figure 6C) [35–37]. Their peptidyl moiety mimicked the sequence around the natural nsp4/5 cut site [17]. To simplify chemical synthesis by basing it on previous approaches, we used an aspartic acid instead of a glutamine at the P1 site and an alanine instead of a threonine in the P6 site.

All three custom peptidomimetic compounds exhibited sub-micromolar IC_50_ values in the FRET-based assay (Figure 6D). Z-ASAVLD-FMK had the highest IC_50_ (0.26 ± 0.02 μM), followed by Z-SAVLD-FMK (0.02 ± 0.001 μM). Z-AVLD-FMK, the shortest of the peptides, exhibited by far the greatest inhibitory potency with an IC_50_ of less than 1 nM (IC_50_ = 0.8 ± 0.09 nM) which is a substantial improvement over the best commercially available peptidomimetic compound Z-VAD-FMK (IC_50_: 0.16 ± 0.01 μM).

To assess whether the customisation also improved the *in vivo* inhibitory capacity, we tested the best custom nsp5 inhibitor (Z-AVLD-FMK) in the viral infection assay (Figure 7A-B). We determined the EC50 value for Z-AVLD-FMK to be 66.01 ± 7.28 μM. Compared to the commercial Z-VAD-FMK, this is a 2-fold increase in inhibitor potency (Figure 7 and Figure 5). This showed that changing the peptidyl moiety to mimic the nsp5 substrate greatly improved its *in vitro* and cell-based inhibitory effect. Taken together, our work indicates that FMK inhibitors targeting nsp5 could be used as a starting point for the development of effective antiviral drugs against SARS-CoV-2.

**Figure 7:**
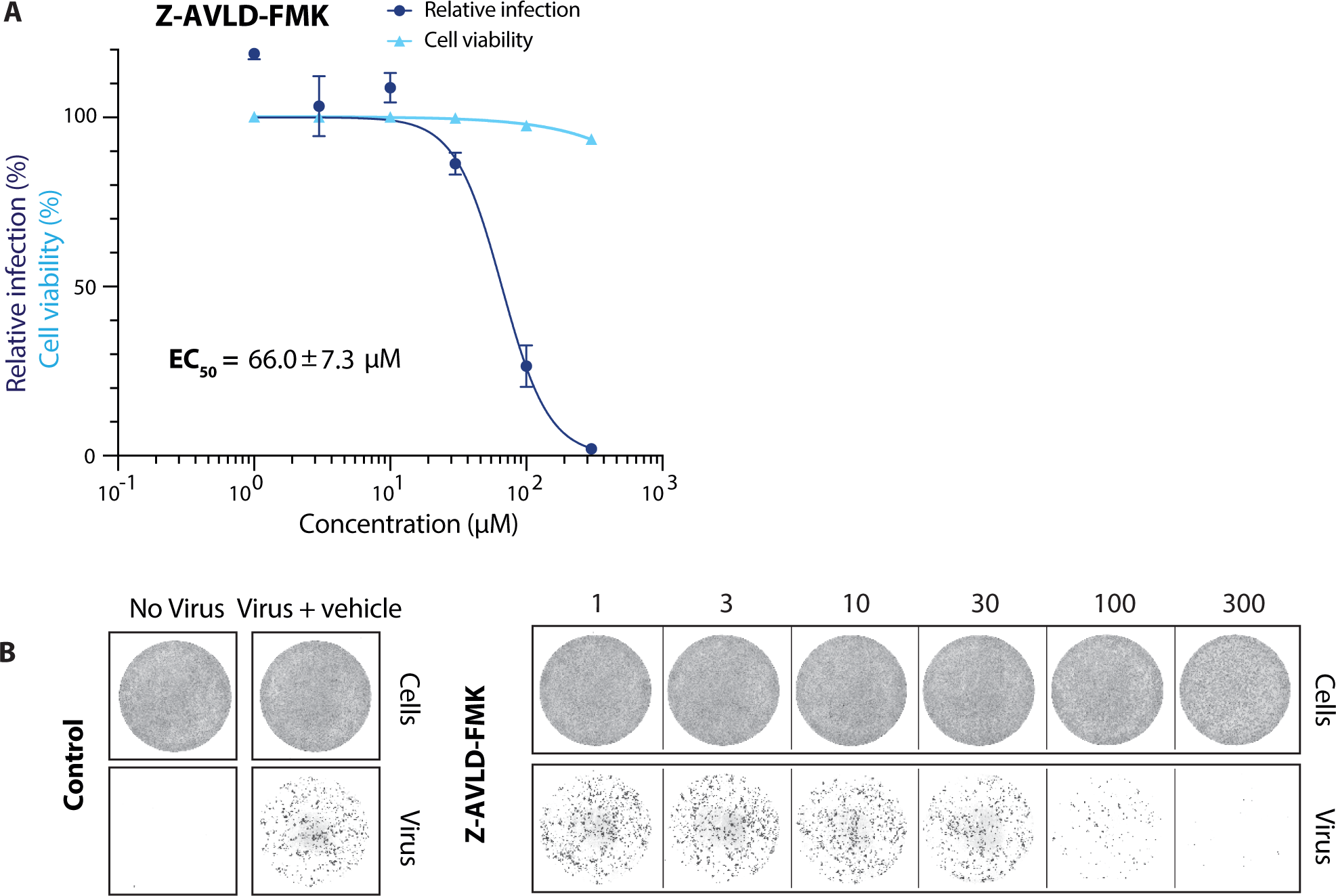
Antiviral activity of improved FMK inhibitor against SARS-CoV-2 in Vero E6 cells. A) Dose response curves for the custom inhibitor Z-AVLD-FMK in a viral infectivity assay in Vero E6 cells. Cell viability values represent the area of cells stained with DRAQ7 DNA dye. Viral infection values were measured as the area of viral plaques visualised by immunofluorescent staining of viral N protein. Data is normalised to DMSO only treated control wells (100 %) and plotted as mean and standard deviation (SD), n=3. Half-maximal effective concentration (EC50) values were determined by non-linear regression. B) Representative images for the SARS-CoV-2 viral infectivity assay in Vero E6 cells, for Z- AVLD-FMK. Representative wells show Vero E6 cells stained for DNA using DRAQ7 (top panel, labelled cells), and viral N protein immunofluorescence (lower panel, labelled virus).

## Discussion

One effective strategy for combatting the COVID-19 pandemic is to re-purpose existing drugs to rapidly identify potential antivirals. The main viral protease is a promising antiviral drug target as it plays a critical role in viral replication by generating the functional viral replication proteins from the polyproteins [13, 14]. Consequently, nsp5 inhibition can halt viral replication and thus might reduce viral load in COVID-19 patients. In our screen of a custom chemical library, we identified calpain inhibitor I and three fluoromethylketone (FMK) peptidomimetics as drugs with antiviral activity against SARS-CoV-2 nsp5 *in vitro* and in a cell-based viral infection assay. In addition, we have further highlighted that the customisation of peptidomimetic FMK inhibitors provide a good starting point for the synthesis of nsp5 specific inhibitors.

Calpain inhibitor I is a synthetic tripeptide aldehyde that acts as a potent inhibitor of cysteine proteases including calpain I, calpain II and cathepsins [38]. It has been tested in mice without obvious toxic effects [39]. Ma *et al* also recently identified Calpain Inhibitor I and several other calpain inhibitors as inhibitors of SARS-CoV-2 nsp5, with IC_50_ values comparable to the results from our HTS in the low μM range [21] in biochemical assays. Our results show that Calpain inhibitor I also inhibited viral growth in a cell-based assay. This suggests that Calpain inhibitors may be useful antiviral drugs against SARS-CoV-2. Three of the strongest hits identified in our HTS were cell permeable FMK peptides. Peptidyl FMKs are widely used due to their ability to strongly and selectively inhibit serine and cysteine proteases, such as caspases, cathepsins and Sentrin/SUMO specific proteases [32, 36]. FMK peptides have been previously investigated as inhibitors of the SARS-CoV-1 nsp5 *in vitro*, and one FMK peptide (Z-DEVD-FMK) has recently been identified as an inhibitor of SARS-CoV-2 nsp5 [24, 40]. The peptidyl backbone of these FMK inhibitors allows them to be utilised for target-based inhibition of specific proteases by mimicking the substrate sequence that binds directly to the active site of the protease. Since the substrate sequence preference of nsp5 is known, we designed three custom inhibitors with the peptidyl moiety based on the nsp5 cleavage site with the predicted highest affinity (nsp4/5) [17]. The shortest of the custom inhibitors, Z-AVLD-FMK, showed extraordinary inhibitor potency *in vitro* with a sub-nanomolar IC_50_. It also improved the inhibitor potency in the cell-based viral infection assay implying that inhibition is a result of the FMK peptides action on nsp5. This shows that peptidomimetic inhibitors are an excellent tool for probing nsp5 protease activity *in vitro* and in cell culture due to their easily customisable nature.

Our custom inhibitors could be further optimised by incorporating a glutamine, found in all natural nsp5 substrate sequences, at the P1 site [8, 18]. Another way forward may be to exchange the FMK functional group with less toxic versions such as difluorophenoxymethylketone (OPh). The development of FMK peptides into clinical drugs was initially halted due to high *in vivo* host cell toxicity. This may be due to the metabolic conversion of the m-FMK group into toxic fluoroacetate, especially in the liver [32]. However, promising studies in mice treated with a peptidyl-OPh inhibitor suggested this alternative functional group it was well tolerated [41]. Furthermore, Q-VD-OPh, has been used in trials investigating AIDS disease progression in rhesus macaques with no toxic side effects [42]. While the commercially available OPh compounds we tested inhibited nsp5 to a lesser extent than the FMK counterparts, customisation of their substrate sequence will likely increase their inhibitory potency against nsp5. Thus, exchanging the functional group of the customised FMK inhibitors could allow them to be developed into clinical drugs.

## Materials and Methods

### Protein purification

#### Nsp5

The nsp5 coding sequence was subcloned from a bacterial expression vector of MBP-nsp5, constructed in MRC PPU (DU67716; available from https://mrcppu-covid.bio/). However, this vector was mis-designed for the C-terminal of nsp5, whereby it only expressed 302 amino acids so the remaining four were added via PCR (forward: 5’ – gaacagattggtggcAGCGGCTTTCGCAAAATGGCG, reverse 5’ – gtgcggccgcttattaCTGAAAGGTCACGCCGCTGCACTGGCGCACCAC). The amplified fragment was then assembled by Gibson assembly in the K27Sumo bacterial expression vector [43] to express 14His-SUMO (N-ter) tagged-nsp5 (306 aa). The final construct (SARS-CoV-2 His-SUMO-nsp5) was then transformed into *E. coli* Rosetta competent cells (Sigma-Aldrich Cat No.71400). Cultures were grown under selective conditions (Kanamycin 50 μg/mL + chloramphenicol 34 μg/mL) at 37 °C until OD 0.8 (600 nm) and expression induced by the addition of 0.5 mM IPTG overnight at 16 °C. Cells were then harvested at 5000 x g for 10 minutes and resuspended in lysis buffer; 50 mM Tris-HCl pH 7.5, 10% glycerol, 1 mM DTT, 500 mM NaCl, 30 mM imidazole and 0.05% NP-40. Cell lysis was done with the addition of 50 μg /mL lysozyme for 10 minutes on ice. Cells were then sonicated at 50% amplitude for 3 x 15s before clarification at 15,000 rpm at 4 °C for 30 minutes in JA-30.50 rotor.

Ni-NTA beads (Thermo Cat. No R90115) were equilibrated in lysis buffer and incubated with crude cell extract for 1 hour at 4 °C. Beads were then collected and washed with lysis buffer before an ATP wash in lysis buffer (+2 mM ATP and 10 mM Mg(OAc)_2_). Samples were then eluted in elution buffer; 50 mM Tris-HCl pH 7.5, 10% glycerol, 1 mM DTT, 500 mM NaCl, 400 mM imidazole and 0.05% NP-40. Nsp5 positive fractions were pooled and incubated with 2 μg/mL of Ulp1 protease overnight at 4 °C to cleave off the 14His-SUMO tag. After cleavage, sample was diluted to 100 mM NaCl in buffer and purified further by anion exchange chromatography over a HiTrap Q column (GE Healthcare (Sigma) Cat. No GE29- 0513-25). The flow through fraction was collected, concentrated to less than 500 μl in a 10K cut off Ultrafiltration Unit (Sigma Cat. No. UFC801024) for a final step of gel filtration over a Superdex 75 column (Sigma-Aldrich Cat. No. GE17-5174-01) into a final buffer; 25 mM HEPES-KOH pH 7.6, 10% glycerol, 0.02% NP-40, 150 mM NaCl and 2 mM DTT.

#### Nsp9 (for gel-based protease assay)

3xFlag-His_6_-SAVLQ-nsp9 (3FH-nsp9) was expressed in baculovirus-infected Sf9 insect cells. The coding sequence of SARS-CoV-2 nsp9 (NCBI reference sequence NC_045512.2) was codon-optimised for *S. frugiperda* and synthesised (GeneArt, Thermo Fisher Scientific). Nsp9 was subcloned into the biGBac vector pLIB [44] to include an N-terminal 3xFlag-His6 tag followed by the last five amino acids of nsp4 (sequence: MDYKDHDGDYKDHDIDYKDDDDKHHHHHSAVLQ-nsp9) (final construct name: SARS-CoV-2 3xFlag-His-nsp5CS-nsp9). Baculoviruses were generated in Sf9 cells (Thermo Fisher Scientific) using the EMBacY baculoviral genome [45]. For protein expression Sf9 cells were infected with baculovirus. Cells were collected 48 h after infection, flash-frozen and stored at -80 °C.

Cell pellets were resuspended in nsp9 buffer (30 mM HEPES pH 7.6, 250 mM sodium chloride, 5 mM magnesium acetate, 10% glycerol, 0.02% NP-40 substitute, 1 mM DTT) supplemented with protease inhibitors (Roche Complete Ultra tablets, 1 mM AEBSF, 10 μg/ml pepstatin A, 10 μg/ml leupeptin) and lysed with a dounce homogenizer. The protein was purified from the cleared lysate by affinity to Anti-FLAG M2 Affinity gel (Sigma-Aldrich) and eluted with nsp9 buffer containing 0.1 mg/ml 3xFlag peptide.

### Gel based nsp9 cleavage assay

Reactions were carried out in a buffer containing 50 mM HEPES-KOH pH 7.6, 1 mM EDTA, 2 mM DTT, 10% glycerol, and 0.02% Tween-20. A concentration of 1 μM nsp5 was used unless otherwise specified. This was incubated with drug compounds resuspended in DMSO or DMSO alone for 10 minutes at room temperature. Reactions were then initiated by the addition of 6.25 μM FLAG-His-SAVLQ-nsp9 substrate. Protease activity was allowed to continue for 1 hour at room temperature before reactions were quenched and denatured by the addition of SDS loading buffer. Products were then separated over a 12% Bis-Tris gel in MOPS buffer and stained with coomassie blue (Generon Cat. No NB-45-00078-1L).

### FRET peptide substrate synthesis

The assay uses a FRET peptide substrate (2-aminobenzoyl – AVLQSGFRKK – tyrosine(3- NO2)R-COOH) which was synthesised in house. The peptides were synthesised on an Intavis ResPep SLi Peptide Synthesiser (CEM, Buckingham, UK) on pre-loaded LL Wang resin using N(a)-Fmoc amino acids including Fmoc-Abz-OH, Fmoc-Tyr(3-NO2)-OH and Fmoc-Gln(trt)-Ser(psi(Me,Me)pro)-OH using HATU as the coupling reagent. Following amino acid chain assembly, peptides were cleaved from the resin by addition to cleavage cocktail (10 mL, 95% TFA, 2.5% H2O, 2.5% TIS) for 2 hours. Following resin removal, peptide precipitation and extensive washing with ether, the peptides were freeze dried. Aliquots of the peptide were dissolved in water and purified on a C8 reverse phase HPLC column (Agilent PrepHT Zorbax 300SB-C8, 21.2x250 mm, 7 m) using a linear solvent gradient of 10- 50% MeCN (0.08% TFA) in H2O (0.08% TFA) over 40 minutes at a flow rate of 8 mL/min.

The peptides were analysed by LC–MS on an Agilent 1100 LC-MSD. The FRET peptide substrate (2-aminobenzoyl- LGARGHRPYD- tyrosine(3-NO2)R-COOH) used with the thrombin FRET protease assay was synthesised by the same method.

### FRET-based nsp5 activity assay

A FRET-based fluorescence quenching assay was developed to monitor hydrolysis activity of nsp5. Upon cleavage of the substrate 2-Aminobenzoyl is no longer quenched by N-Tyrosine and able to fluorescence. Any changes in fluorescence were monitored in the excitation spectra of 330 nm and emission spectra of 420 nm. Unless otherwise stated, the reaction master mix used in all FRET experiments contributed to a final concentration of; 10 nM nsp5, 50 mM HEPES-KOH pH 7.6, 1 mM EDTA, 2 mM DTT, 10% glycerol, and 0.02% Tween. Reactions were initiated by the addition of FRET substrate to a final concentration of 20 µM (unless otherwise stated) to reaction master mix. A Tecan M1000 pro or Tecan Spark were used to take fluorescence readings for all FRET experiments with the following settings: Kinetic cycle: 10, Interval time: 2 minutes, Excitation bandwidth: 5nm, Emission bandwidth: 15 nm, Gain: 140, Number of flashes: 50, Flash frequency: 400 Hz, Integration time: 20 µs and Z-position: 22470 µm. All nsp5 protease reactions were done at room temperature (19°C - 23°C) however, ambient temperature inside the Tecan M1000 pro or Tecan Spark plate readers fluctuated between 25°C and 27.5°C.

### K_m_ determination of nsp5 for FRET-based substrate

To determine the K_M_ for our FRET substrate, the FRET-based nsp5 activity assay was done as described but with a substrate concentration range of 0 – 75 μM. Initial velocities of the proteolytic activity were calculated by linear regression of the first 14 minutes of the kinetic progress curve and K_M_ values were calculated by fitting to a Michaelis-Menten equation using GraphPad PRISM (Version 8.4.3). These data were corrected for inner filter effect by the method described by Liu et. al [46].

### Nsp5 High throughput screen (HTS)

To identify inhibitors of nsp5 the FRET-based nsp5 activity assay, as described above, was used to screen a custom library of over 5,000 compounds assembled from different sources (Sigma, Selleck, Enzo, Tocris, Calbiochem, and Symansis). All drug compounds present in the custom library were stored at a stock concentration of 10 mM in DMSO. Two concentrations of the custom library were used in the HTS. In the high concentration (4 µM) 12.5 nL of compound was distributed into 24 384-well plates (Greiner 384 Flat Bottom Black Polystyrene Cat. No.: 781076) using an Echo 550 (Labcyte) onto 1µL DMSO into columns 3-22. At the lower concentration (0.8 µM) 2.5 nL of drug compound was distributed onto 1 µL DMSO in the same way. Each HTS plate had a total of 4 columns (1-2, 23-24) without drug compounds which were used for control reactions. Columns 1 and 23 contained a positive control of enzyme with no drug and then columns 2 and 24 served as negative controls of reactions with no enzyme. After reaction master mix (22.5 µL, as described above) was dispensed into 384-well reaction plates by XRD-384 Automated Reagent Dispensers (FluidX Ltd). Plates were then centrifuged briefly at 1,000 x g and incubated at room temperature for 10 min. To start the reaction, 12.5 µl of 20 µM FRET substrate was dispensed and the plates were spun briefly. Following reaction initiation, fluorescence readings were taken every 2 min for a total of 10 time points from 1 minute 20 seconds into the reaction.

### Identification of Primary Hits

Time-course data was analysed using MATLAB (R2020_a) to determine the rate of reaction then normalised to the rate of reaction of positive (no drug compound) control reaction wells. We assessed each individual HTS screen plate and the HTS as a whole by calculating a Z’ factor score based on normalised rate of reaction compared to no enzyme controls. The screen had a Z’ factor greater than 0.6 indicating that we could reliably identify hits. Percentage reduction in rate of reaction or “activity” were then calculated for each drug compound. Those with a reduction in activity that was greater than 30% in the highest drug concentration (4 µM) were selected as primary hits. Hits were then ranked based on percentage reduction in activity and whether they were hits in both HTS concentrations.

### Compound resuspension for hit validation

Compounds selected for *in vitro* experimental validation and cell-based experiments were purchased and resuspended into DMSO following manufacturer’s instruction.

### Half maximal inhibitory concentration (IC_50_) determination

Drug compounds for IC_50_ determination were titrated over a wide range from 1500 mM to either 1 nM over 20 steps or from 500 mM to 10 pM over 30 steps depending on the drug stock concentrations available. FRET-based nsp5 activity assays were carried out, as described above. The rate of the reaction was determined from the time-course data and normalised to positive control (no drug) samples. IC_50_ values were then calculated using non-linear regression of inhibitor concentration against the normalised response using variable slope via GraphPad Prism (Version 8.4.3) software.

### Viral inhibition assay

SARS-CoV-2- isolate and recombinant antibodies against viral nucleocapsid protein used for the viral infection assay were produced in-house as described elsewhere (Zheng et al, Biochem J, this issue). The conditions for the viral inhibition assay in Vero E6 cells are described in Zheng et al (Biochem J, this issue).

## Data Availability Statement

All data associated with this paper will be deposited in FigShare (https://figshare.com/).

## Author Contributions

**Jennifer C Milligan & Theresa U Zeisner:** Conceptualization, methodology, investigation, software, validation, formal analysis, data curation, writing – original draft, Writing - review & editing, visualization. **George Papageorgiou:** Methodology, investigation, data curation, writing - review & editing. **Dhira Joshi:** Methodology, investigation, data curation. **Christelle Soudy:** Methodology. **Rachel Ulferts:** Methodology, investigation. **Mary Wu:** Methodology, resources, investigation. **Chew Theng Lim:** Methodology, investigation. **Kang Wei Tan:** Methodology, investigation. **Florian Weissmann:** Methodology, investigation, Writing - review & editing. **Berta Canal:** Methodology, investigation, Writing - review & editing. **Ryo Fujisawa:** Methodology, investigation. **Tom Deegan:** Methodology, investigation. **Hema Nagara:** Resources. **Ganka Bineva-Todd:** Resources. **Clovis Basier:** Investigation. **Joseph F Curran:** Investigation, writing - review & editing. **Michael Howell:** Supervision, resources, project administration, funding acquisition. **Rupert Beale:** Supervision, project administration. **Karim Labib:** Supervision, project administration, writing - review & editing. **Nicola O’Reilly:** Supervision, resources, project administration. **John F.X Diffley:** Conceptualization, supervision, methodology, project administration, funding acquisition, writing - review & editing.

## Acknowledgments

We thank all members of the involved labs for their help and comments on the manuscript. We thank Peptide Chemistry (STP) and High Throughput Screening (STP) at the Francis Crick Institute for all their help and are grateful to MRC Reagents and Services (https://mrcppureagents.dundee.ac.uk/) for providing DNA constructs for nsp5. This work was supported by the Francis Crick Institute, which receives its core funding from Cancer Research UK (FC001066), the UK Medical Research Council (FC001066), and the Wellcome Trust (FC001066). This work was also funded by a Wellcome Trust Senior Investigator Award 106252/Z/14/Z to JFXD; the UK Medical Research Council MC_UU_12016/13 and Cancer Research UK Programme Grant C578/A24558 to KL; a Sir Henry Wellcome Postdoctoral Fellowship 204678/Z/16/Z to TD). TUZ received funding from the Boehringer Ingelheim Fonds. BC and FW have received funding from the European Union’s Horizon 2020 research and innovation programme under the Marie Skłodowska- Curie grant agreement Nos 895786 and 844211.

## Supplemental Material

**Figure S1:**
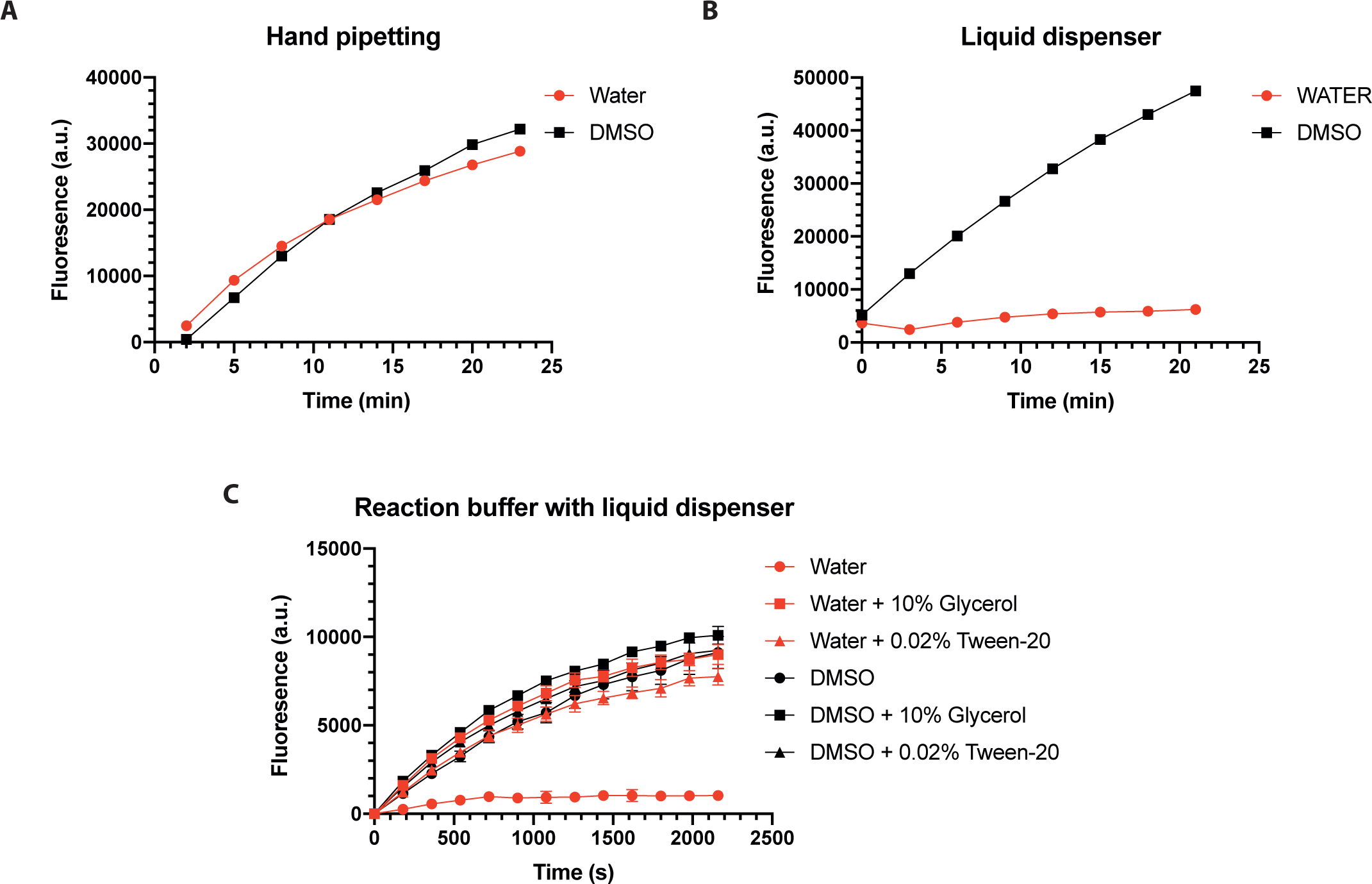
Testing reaction buffers for high-throughput screening. A - B) Increase in fluorescence over time after initiation of nsp5 protease FRET-based assay onto either DMSO or water via hand pipetting (A) or via an automated liquid handling robot (B). Experiments were done in triplicate. C) The nsp5 protease FRET-based assay was monitored for increase in fluorescence over time in standard buffer (50 mM Hepes-KOH pH 7.6, 2 mM DTT and 1 mM EDTA) onto either DMSO or water. 10% glycerol or 0.02% Tween are added to the standard reaction buffer which is dispensed onto either DMSO or water.

**Figure S2:**
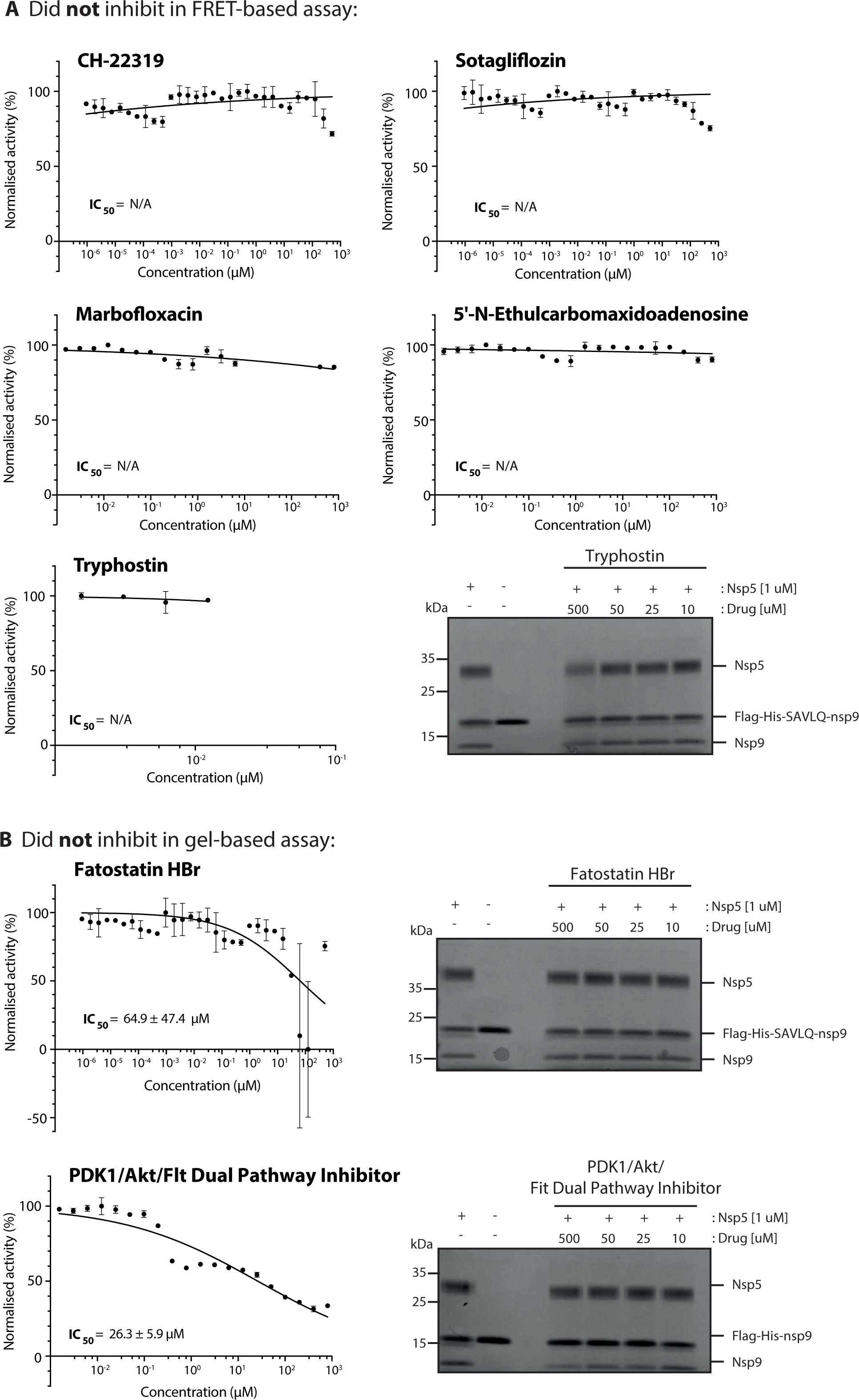
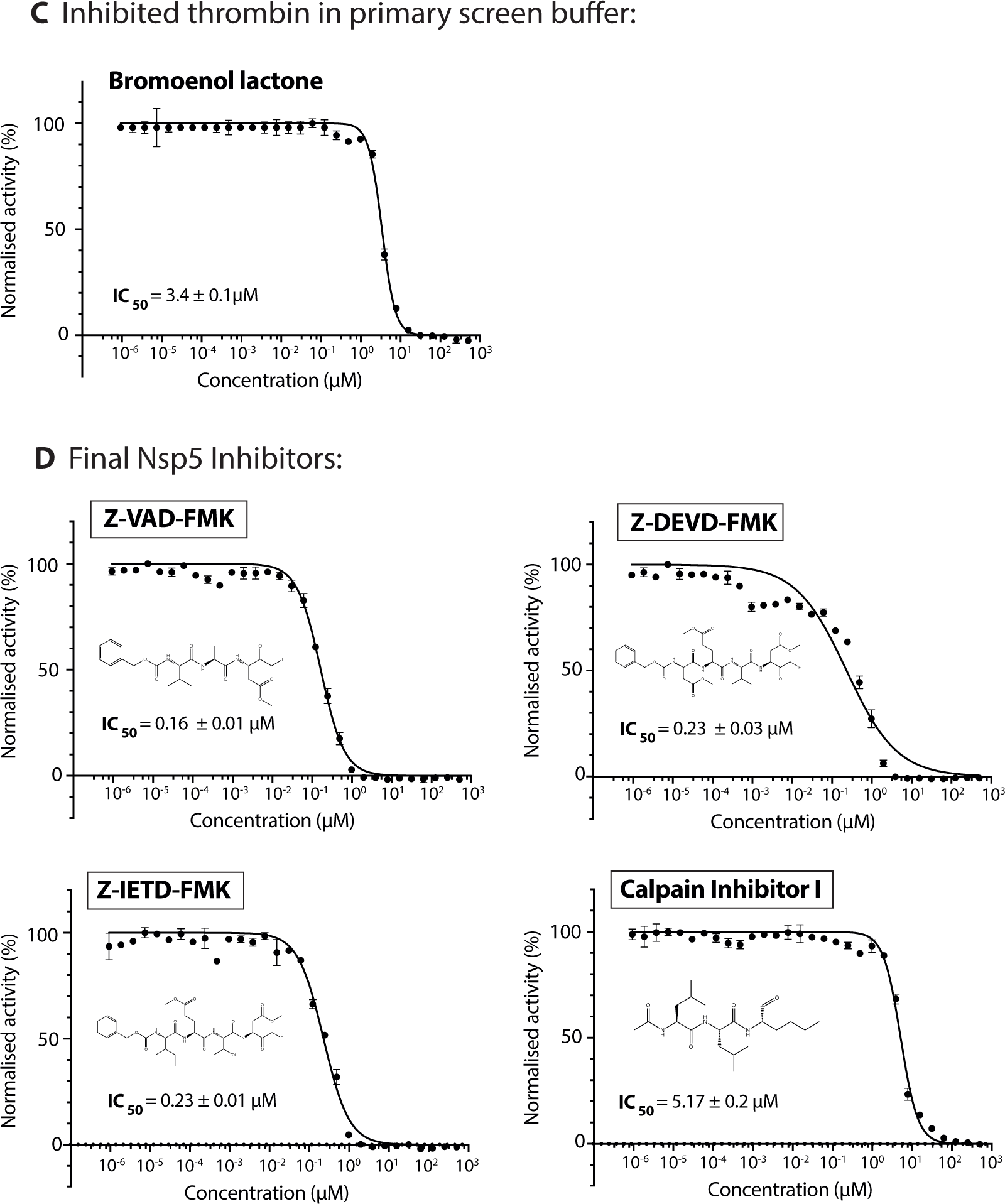
IC_50_ values and related figures of all hits. A-D) *In vitro* activity of nsp5 was measured over a wide-range concentrations of different inhibitors using the FRET-based assay. A dose response curve for an IC_50_ value was determined by non-linear regression (shown in black) for all drugs where possible. Drugs are sorted into different stages of validation. All FRET-based data are shown as mean ± SEM, n=3, error bars represent SD. For select compounds a nsp9 gel-based assay was done at a wide range of drug concentrations. (D) Calpain Inhibitor I, Z-VAD-FMK, Z-DEVD-FMK and Z-IETD-FMK were all validated as specific nsp5 inhibitors. FRET-based assay dose dependent curves are shown with IC_50_ values determined by non-linear regression with normalised data alongside relevant chemical structures. All data are shown as mean ± SD, n=3, error bars represent SD.

**Figure S3:**
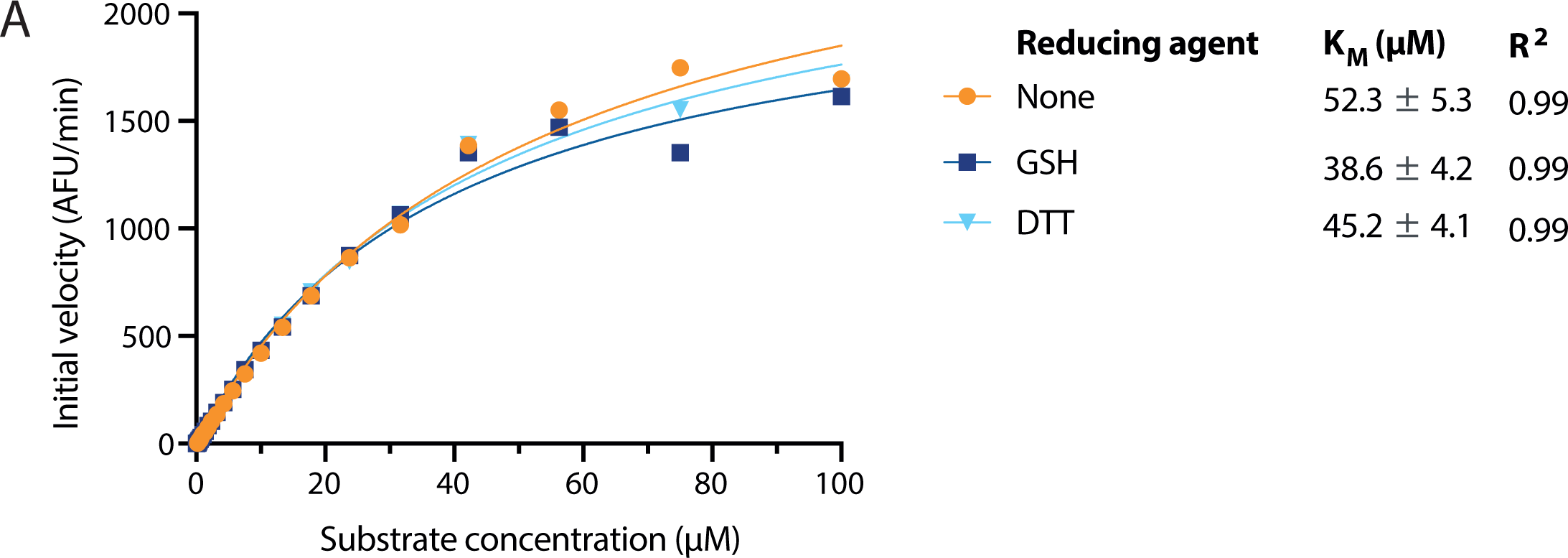
Enzyme characterisation in different reducing environments. A) Graph of K_M_ determination. Initial reaction rates at a range of concentrations were plotted against substrate concentration to obtain values for K_M_ using different reducing reagents in the buffer (no reducing agent, DTT, GSH).

**Figure S4:**
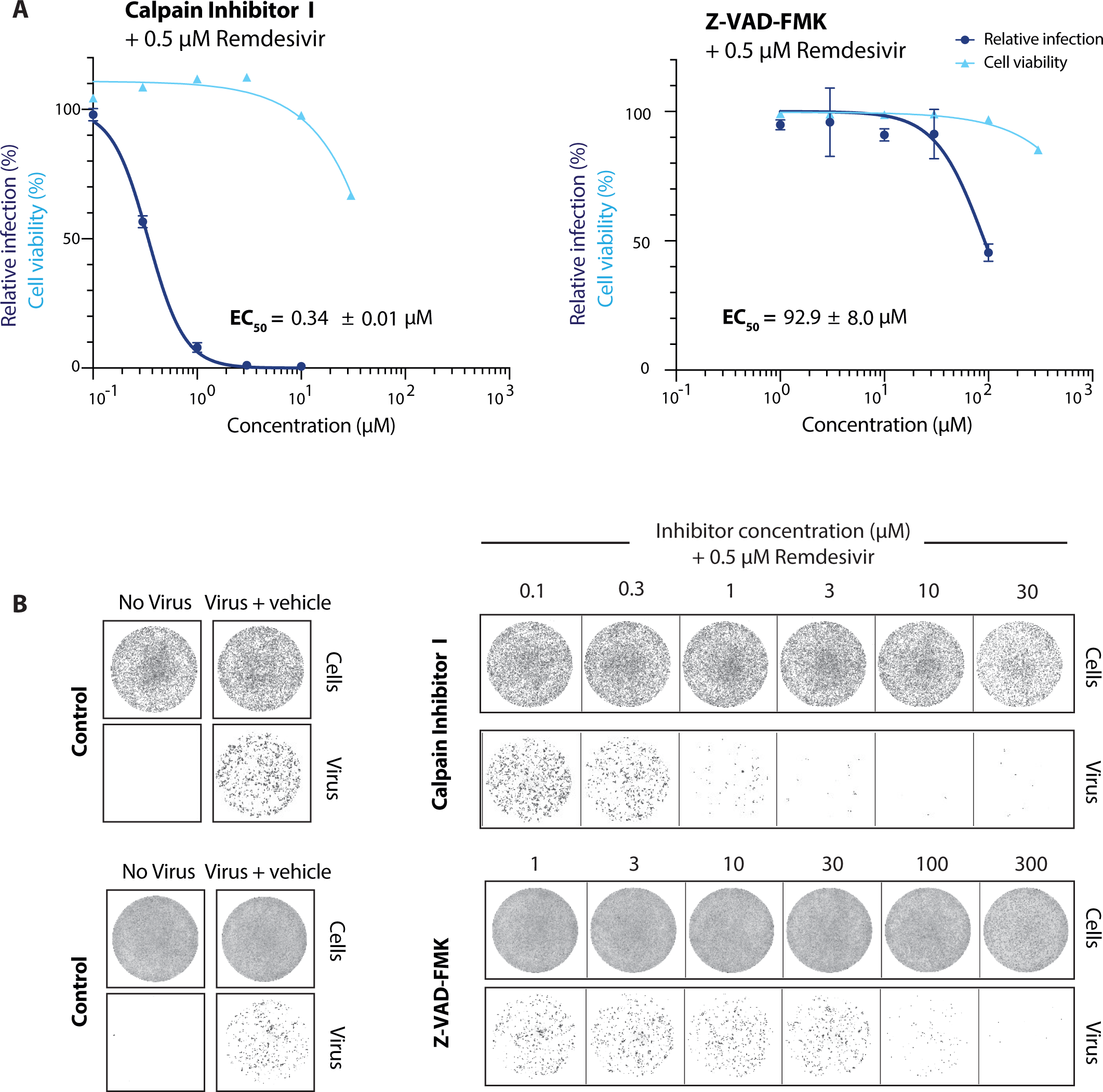
Antiviral activity of HTS hits in combination with remdesivir against SARS-CoV-2 in Vero E6 cells. A) Dose response curves for the validated hits (Calpain Inhibitor I and Z-VAD-FMK) in combination with remdesivir in a viral infectivity assay in Vero E6 cells. Cell viability values represent the area of cells stained with DRAQ7 DNA dye. Viral infection values were measured as the area of viral plaques visualised by immunofluorescent staining of viral nucleocapsid protein. Data is normalised to DMSO only treated control wells (100 %) and plotted as mean and standard deviation (SD), n=3. Half-maximal effective concentration (EC50) values were determined by non-linear regression. B) Representative images for the SARS-CoV-2 viral infectivity assay in Vero E6 cells, for Calpain Inhibitor I and Z-VAD-FMK in combination with remdesivir. Representative wells show Vero E6 cells stained for DNA using DRAQ7 (top panel, labelled cells), and viral N protein immunofluorescence (lower panel, labelled virus).

**Figure S5:**
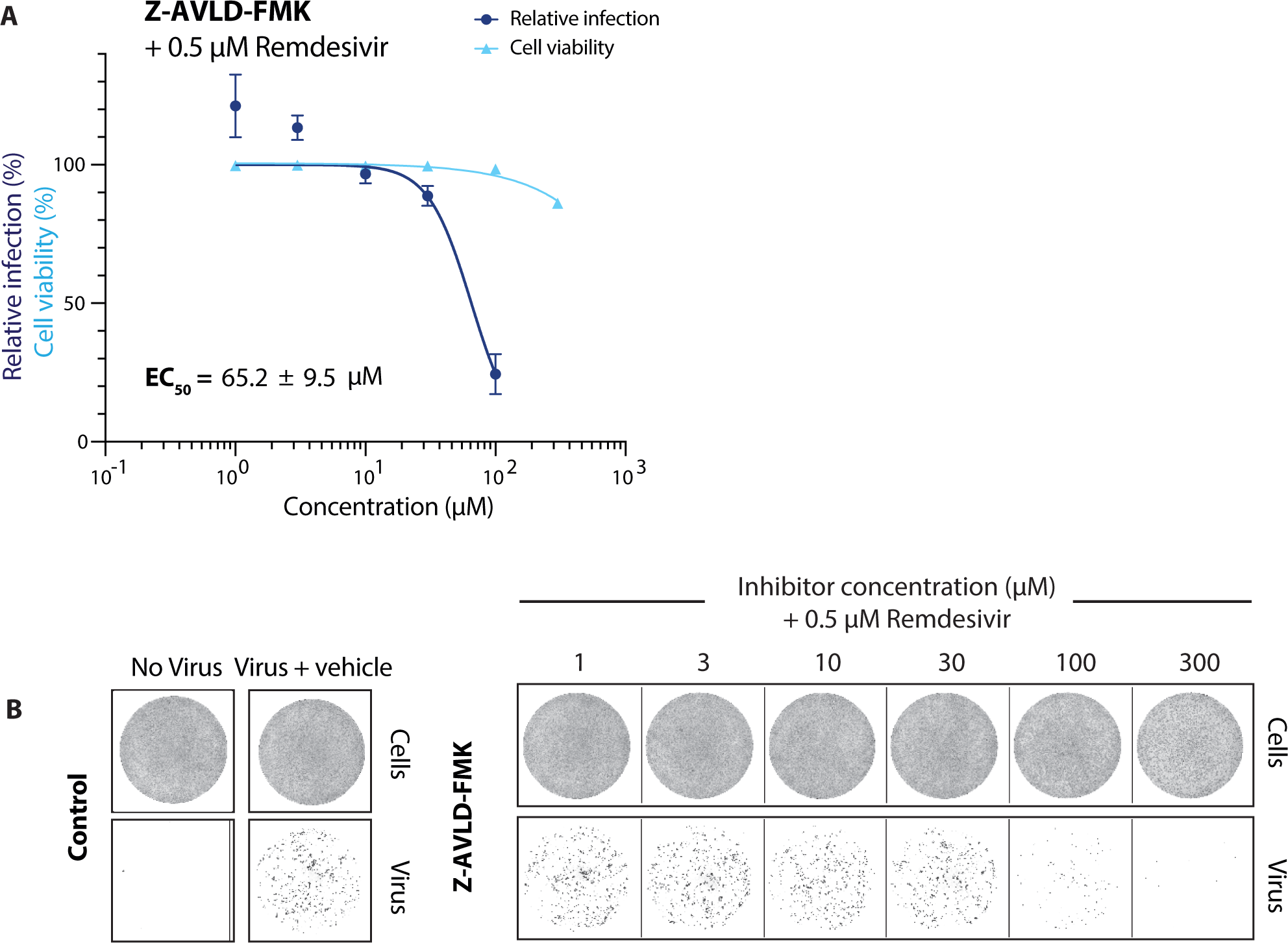
Antiviral activity of improved FMK inhibitor in combination with remdesivir against SARS-CoV-2 in Vero E6 cells. A) Dose response curves for the custom inhibitor Z-AVLD-FMK in combination with remdesivir in a viral infectivity assay in Vero E6 cells. Cell viability values represent the area of cells stained with DRAQ7 DNA dye. Viral infection values were measured as the area of viral plaques visualised by immunofluorescent staining of viral nucleocapsid protein. Data is normalised to DMSO only treated control wells (100%) and plotted as mean and standard deviation (SD), n=3. Half-maximal effective concentration (EC50) values were determined by non-linear regression. B) Representative images for the SARS-CoV-2 viral infectivity assay in Vero E6 cells, for Z- AVLD-FMK in combination with remdesivir. Representative wells show Vero E6 cells stained for DNA using DRAQ7 (top panel, labelled cells), and viral N protein immunofluorescence (lower panel, labelled virus).

